# Green leaf volatiles co-opt proteins involved in molecular pattern signaling in plant cells

**DOI:** 10.1101/2022.10.15.512371

**Authors:** Sasimonthakan Tanarsuwongkul, Kirsten Fisher, B. Todd Mullis, Jamie Roberts, Harshita Negi, Qiang Wang, Johannes W. Stratmann

## Abstract

The green leaf volatiles (GLVs) *Z*-3-hexen-1-ol and *Z*-3-hexenyl acetate are airborne infochemicals released from damaged plant tissues that prime defenses against herbivores and pathogens in receiver plants. They are conceptually similar to well-known damage-associated molecular patterns (DAMPs), but little is known about their mechanism of action. Using tomato cell cultures, we found that rapid responses to the two GLVs and the polypeptide DAMP systemin showed a significant overlap but also GLV-specific patterns. Within five minutes, GLVs induced changes in MAPK activity and proton-fluxes as well as rapid and massive changes in the phosphorylation status of proteins. Many of these proteins are involved in reprogramming the proteome from cellular homeostasis to stress and include pattern recognition receptors, a receptor-like cytoplasmic kinase, MAPK cascade components, calcium signaling proteins, and transcriptional regulators, all of which are also components of DAMP signaling pathways. This phosphoproteome may represent an early priming state that enables plants to respond forcefully to a subsequent stress signal.

## Introduction

Green leaf volatiles (GLVs) are small volatile organic compounds (VOCs) that are synthesized from fatty acids derived from chloroplast membrane lipids. GLVs are released within seconds to minutes in response to tissue damage by wounding or herbivory as well as pathogen infections and abiotic forms of stress ^1–3^. As airborne infochemicals they reach undamaged parts of a damaged plant or neighboring receiver plants and thus enable intra- and interplant communication. GLVs are released from damaged cells and induce defenses, similar as damage-associated molecular patterns (DAMPs) ^4–7^. GLV perception results in upregulation of defenses or priming in receiver tissues ^8,9^. Priming does not result in upregulation of a full-fledged defense response, which is resource-intensive ^10^. In the absence of an attacker, relatively few resources are dedicated to defenses during priming. In contrast, if an attack is directed towards a primed plant, defenses can be deployed faster and stronger than in unprimed plants ^11^. A primed state is established through transcriptional and metabolic reprogramming in response to various GLVs in many angiosperm plants ^12^.

While the role of GLVs in priming is well established, it is unknown how the GLV signal is perceived by plant cells, and little is known about GLV-induced signal transduction when compared to well-characterized elicitors of plant defenses such as microbe-, damage-, and herbivore-associated molecular patterns (MAMPs, DAMPs, and HAMPs). These molecular patterns are structurally diverse but not similar to GLVs and not volatile. They are perceived by plasma membrane-bound pattern recognition receptors (PRRs), which consist of an extracellular ligand-binding domain, a transmembrane domain, and an intracellular kinase domain ^13^. Animals possess odorant binding proteins that deliver VOCs to olfactory receptors ^14^, but to our knowledge, no plant homologs of these receptors ^15^ and no plant odorant binding proteins have been discovered. Because GLVs can pass membranes either passively or via transporters ^16,17^, they may exert their effects through interaction with intracellular binding proteins. However, there are multiple membrane-diffusible ligands that are perceived by plasma membrane-localized receptors including brassinosteroids by BRI1 ^18^, hydrogen peroxide by HPCA1^19^, medium chain 3-hydroxy fatty acids by LORE ^20^, a methyl sphingoid by RDA2 ^21^, and the volatile plant hormone ethylene by two-component His protein kinase receptors ^22^.

There are very few structure-function analyses for GLVs. Heil et al. ^23^ showed that there were no significant differences among (*Z*)-3-hexenyl acetate (Z3-HAC), E3-HAC, E2-HAC, 5-HAC, (*Z*)-3-hexenyl isovalerate and (*Z*)-3-hexenyl butyrate regarding their potential to induce extrafloral nectar secretion in lima beans, an indirect defense response. This raises the question as to whether these structurally diverse GLVs with different chemical properties are all perceived by the same mechanism and activate a common signaling pathway that induces nectar secretion. Other ideas for GLV/VOC perception are that some hydrophobic VOCs could alter membrane properties such as permeability for ions that may result in membrane depolarizations ^23,24^, or that VOCs induce a change in the cellular redox potential and thus redox-regulated responses ^25^. However, these unspecific mechanisms are hard to reconcile with the activation of relatively specific transcriptional and defensive responses such as extrafloral nectar production.

Only a few VOC-induced signaling components are known, and even fewer GLV-specific ones. *E*-2-Hexen-1-al (E2-HAL), Z3-HAL, Z3-HAC and *E*-2-hexen-1-ol (E2-HOL) induced rapid increases in cytosolic Ca^2+^ concentrations in Arabidopsis and tomato, and E2-HAL, as a reactive carbonyl, resulted in the generation of the reactive oxygen species (ROS) superoxide anion, whereas E2-HOL was inactive ^24,26^. In addition, these GLVs caused a rapid membrane potential depolarization, which can be associated with calcium and ROS signaling ^27^. Furthermore, GLVs from various sources induced phosphorylation of two mitogen-activated protein kinases (MAPKs) in the grass *L. temulentum* ^28,29^. MAPK cascades are common phosphorylation-dependent signaling modules in many stress signaling pathways. Calcium-, ROS-, and MAPK-signaling is also rapidly induced by molecular patterns and involved in the synthesis of stress hormones such as salicylic acid (SA) and jasmonic acid (JA) ^30–33^. Z3-HAC, Z3-HOL, and Z3-HAL induced JA synthesis in maize ^11,34^, which may be part of the mechanism underlying priming in maize, and also in *N. benthamiana* and tomato ^11,35,36^. In line with these findings, transcriptomics analyses in a range of species corroborate a prominent role of JA in responses to GLVs, and also identified GLV-responsive genes involved in Ca^2+^- and MAPK signaling, as well as defense genes involved in direct defenses against herbivores and microbial pathogens ^37 38 39^. In addition, a forward genetics approach in Arabidopsis identified mutants that do not respond to E2-HAL. The mutations were found in a gamma-aminobutyric acid (GABA) transaminase gene that codes for an enzyme that degrades GABA ^40^, and in an oxidoreductase ^41^. GABA is also involved in the activation of defenses against pathogens ^42^.

In summary, while very little is known about GLV perception, some signal transduction processes and some metabolic events with implications for GLV-signaling have been identified. However, there is no coherent study that delineates a GLV-induced signaling network as known for MAMPs and DAMPs. Studies were carried out with different GLVs, different plant systems, and under different conditions. Therefore, it is unknown how a specific defense response, such as priming against herbivores, is established and what constitutes priming by GLVs at the molecular level.

In this work, using suspension-cultured cells of the wild tomato species *Solanum peruvianum* (SP cells), we identified Z3-HOL- and Z3-HAC-induced cellular responses and phosphoproteins that may function in GLV-induced signaling pathways (Fig. 1). These proteins include receptors and signaling proteins that also operate in classical molecular pattern signaling pathways, as determined by including the DAMP systemin in our analysis. Since the observed signaling events occurred within five minutes of GLV exposure, they may represent very early molecular mechanisms that establish a state of priming.

**Fig. 1.**
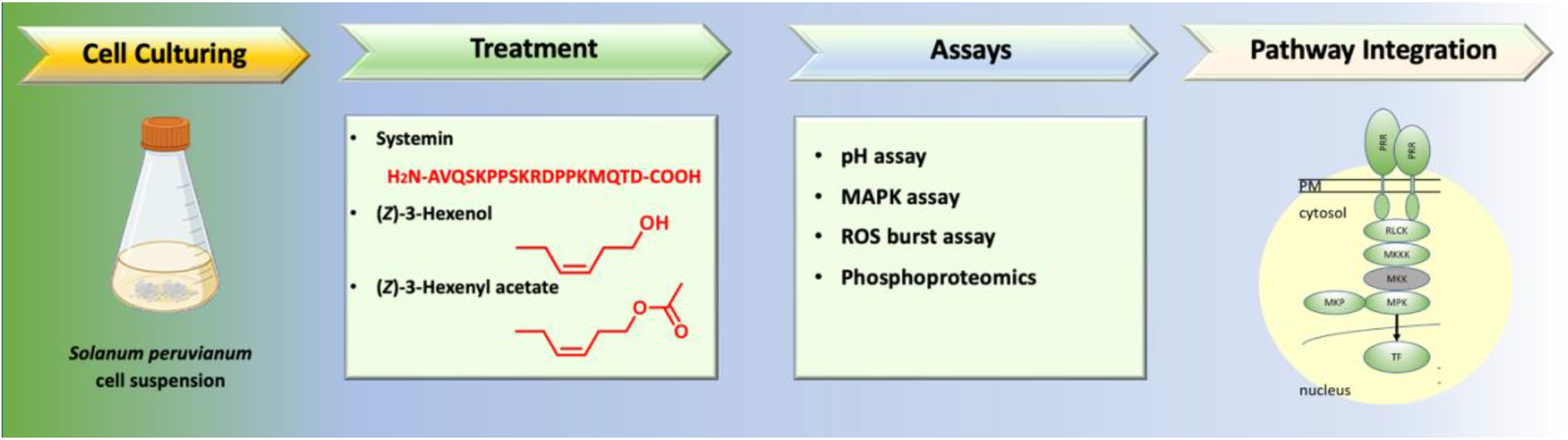
Experimental flow chart. SP cells were treated with GLVs and systemin, followed by testing of rapid cellular responses. Identified response proteins were assigned pathway positions.

## RESULTS

### Z3-HOL and Z3-HAC induce a change in the pH of cultured tomato cells

A well-known early PRR-mediated signaling response is the alkalinization of the growth medium of suspension-cultured plant cells ^43–45^. We found that the GLVs Z3-HOL and Z3-HAC, which are emitted by tomato plants in response to herbivory and JA-treatment ^2^, also induce a medium alkalinization response in SP cells. The two GLVs induced distinct pH response profiles in a concentration-dependent manner while the solvent control ethanol was inactive. The response to Z3-HOL most closely resembled the response to systemin, although the lowest concentration of Z3-HOL that induced an alkalinization response was much higher (5 mM) than for systemin (10 nM) (Fig. 2A), probably due to low solubility of Z3-HOL in the medium. The SP cells were more sensitive to Z3-HAC, which caused a medium acidification response at concentrations as low as 270 μM (Fig. 2B). The initial acidification response lasted 20 to 30 min, after which the pH started to increase. This biphasic pH response distinguishes Z3-HAC from Z3-HOL, systemin, and other DAMPs and MAMPs tested on cultured plant cells, which generally induce a simple alkalinization response ^43–45^. We also measured pH changes within the first 5 min after addition of the GLVs and systemin at a high temporal resolution (Supplementary Fig. 1A). This revealed a 1 to 1.5 min lag time before the start of the pH increase or decrease, which is typical for PRR-mediated responses. To exclude that increases in the pH of the culture medium are not a consequence of proton influx due to irreversible damage to the plasma membrane, we showed that SP cells were responsive to the proton ATPase agonist fusicoccin when applied 90 min after Z3-HOL and Z3-HAC treatments, resulting in medium acidification and thus recharging of the proton gradient across the plasma membrane (Supplementary Fig. 1B,C).

**Fig. 2.**
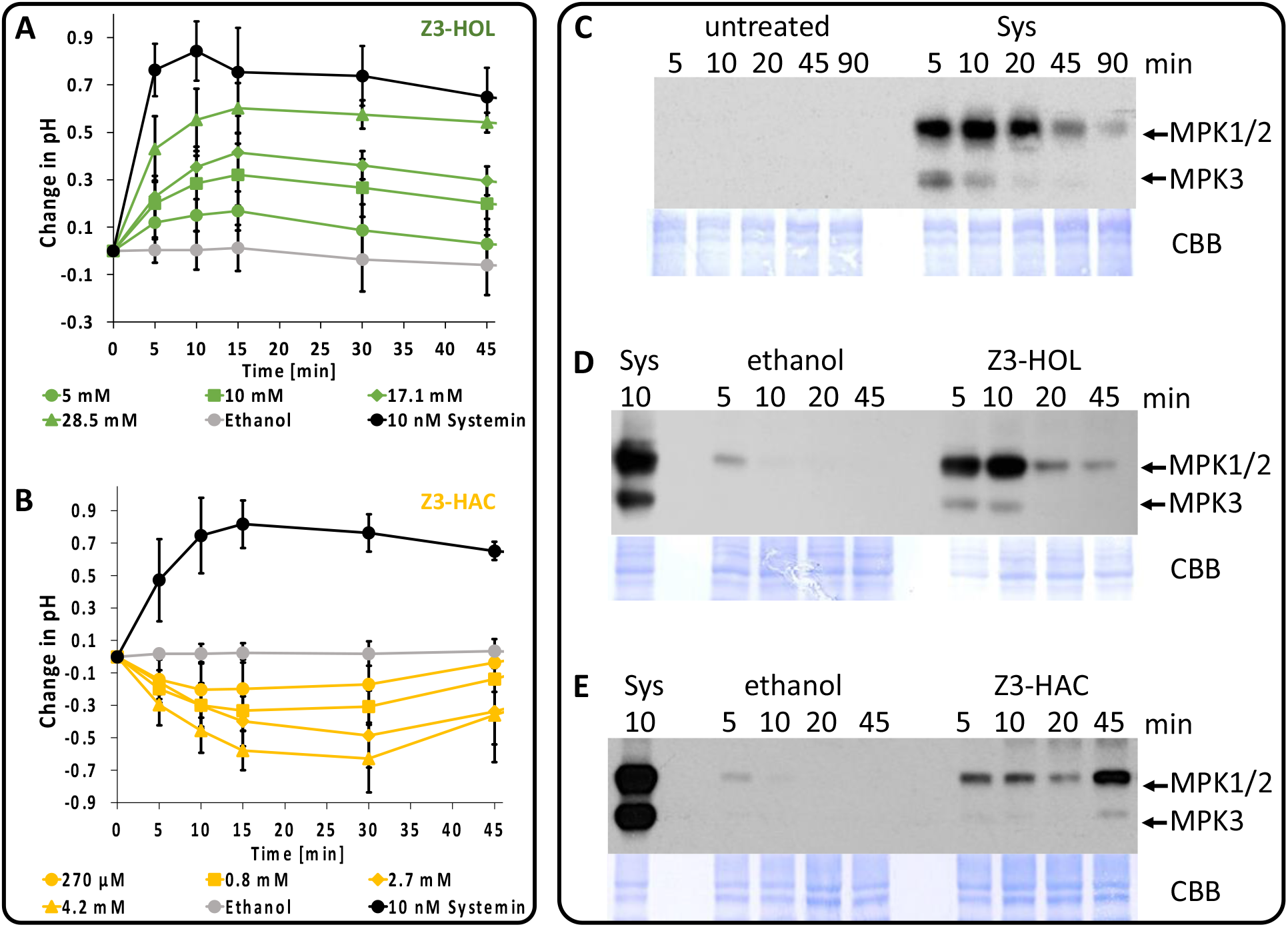
Distinct kinetics of Z3-HOL- and Z3-HAC-induced medium pH changes and MAPK phosphorylation in SP cells. **A,B** For pH assays, SP cells were treated with 10 nM systemin, various concentrations of Z3-HOL (**A**) or Z3-HAC (**B**) solved in 10 μl ethanol, and 10 μl ethanol alone (**A,B**). The medium pH was measured over a period of 45 min. The change in medium pH is expressed as the increase or decrease in the medium pH above the pH at time = 0. Graphs represent the average and standard deviations of four independent experiments (n = 4). **C-E** For MAPK assays, SP cells were treated with 3 nM systemin (**C-E**), 29 mM Z3-HOL (**D**) or 4.2 mM Z3-HAC (**E**) solved in 15 μl ethanol, and 15 μl ethanol alone (**D,E**). Phosphorylation of MAPKs were visualized by immunoblotting using an antibody against the phosphorylated activation motif pTEpY. Lower panels show coomassie brilliant blue (CBB) stained membranes. Sys – systemin. Similar results were obtained in at least three independent experiments (n ≥ 3). Representative experiments are shown.

### Z3-HOL and Z3-HAC induce phosphorylation of MAPKs in cultured tomato cells

Another typical signaling response to molecular patterns is the activation of mitogen-activated protein kinases (MAPKs) via phosphorylation. We had shown earlier that systemin and MAMPs like flg22 and chitin strongly activate the tomato MAPKs MPK1, MPK2, and weakly MPK3 in SP cells ^46,47^. Using an antibody against the phosphorylated activation motif pTEpY of MAPKs we showed that 28.5 mM Z3-HOL and 4.2 mM Z3-HAC, concentrations that induced a strong pH response, also induced MAPK phosphorylation in SP cells. MPK1 and MPK2 evolved via a recent gene duplication and could not be distinguished using this antibody ^47^. Systemin (3 nM) was included as a reference; it induced an early peak of MAPK phosphorylation at 5 min (MPK3) and 10 min (MPK1/2), followed by a continuous decrease (Fig 2C). The two GLVs also induced pronounced MAPK phosphorylation, but with distinct kinetics, whereas the response to ethanol was very weak and transient. Similar to systemin, Z3-HOL induced rapid (within 5 min) phosphorylation of MPK1/2 and MPK3 which strongly decreased after a peak at 10 min (Fig. 2D). In contrast, Z3-HAC induced biphasic MPK1/2 phosphorylation and a late MPK3 response after 20 min, correlating with a biphasic pH response (Fig. 2E).

### GLVs do not induce an oxidative burst in tomato leaf cells

Many MAMPs and DAMPS induce an extracellular oxidative burst in leaf discs of various plant species ^48^. However, it was shown that systemin does not induce such an oxidative burst ^46,49^, unlike other peptide DAMPs such as AtPep1^48^. Z3-HOL and Z3-HAC also did not induce an oxidative burst, although various concentrations were tested. By constrast, the MAMP flg22 caused a pronounced oxidative burst (Supplementary Fig. 2).

### GLV- and systemin-induced rapid phosphorylation profiles

To further investigate early signaling events that are closely tied to GLV perception, we pursued a phosphoproteomics approach to identify rapidly modified signaling proteins. We treated SP cells for five minutes with the two GLVs and systemin at the same concentrations as used for the pH and MAPK assays. Controls were ethanol-treated cells for GLVs and untreated cells for systemin. All results represent three independent experiments. At five minutes after treatment, no changes in protein abundance via translation or degradation are expected, as shown by Wu et al. (2014)^50^.

Using an automated workflow ^51^ and the PolyTi resin for phosphopeptide enrichment, ^51–53^ we identified ~ 1700 phosphoproteins at medium to high confidence. The average specificity of phosphopeptide enrichment was ≥ 79% (Fig. 3A, B). This confirmed that the enrichment method developed by Mullis et al. ^51^ is highly specific toward phosphopeptides. On average, 88.6% of all phosphorylations were on serine, 10.4% on threonine, and 0.9% on tyrosine (Fig. 3C). Most peptides were singly phosphorylated (74.7%) (Fig. 3D).

**Fig. 3.**
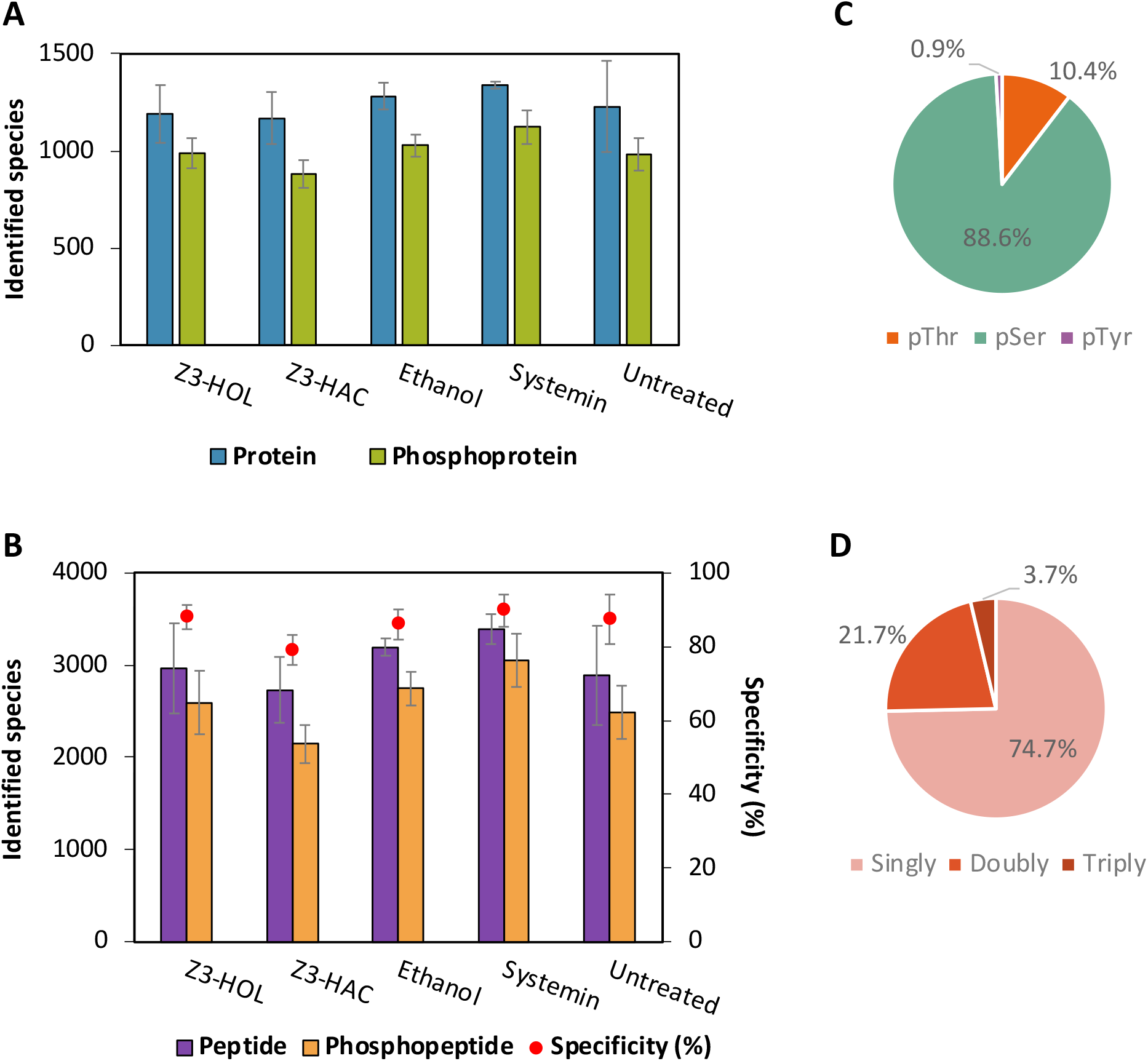
Analyses of label-free quantitative phosphoproteomics using PolyTi resins for phosphopeptide enrichment. **A** The average number of total identified proteins (blue bars) as compared to phosphoproteins (green bars) are shown for each treatment. **B** The average number of total identified peptides (purple bars) as compared to phosphopeptides (orange bars) is shown and phosphopeptide enrichment specificity (red circle), which is the percentage of phosphopeptides among all identified peptides. **C** Frequency of phosphoserine (pSer), phosphothreonine (pThr) and phosphotyrosine (pTyr) residues in the dataset. **D** Distribution of singly, doubly and triply phosphorylated peptides.

Label-free quantitation was performed by calculating the fold change of a treatment compared to controls using Proteome Discoverer. The cutoff for significance for the calculated fold change (abundance ratio) was set at greater than or equal to 2 for phosphorylation and less than or equal to 0.5 for dephosphorylation. This resulted in identification of 282, 184, and 768 phosphoproteins with an abundance ratio of ≥ 2.0 and 64, 281, and 18 phosphoproteins with an abundance ratio of ≤ 0.5 for Z3-HOL, Z3-HAC, and systemin, respectively (Supplementary Table 1). By far the highest number of protein phosphorylations were induced by systemin. Of the GLV-induced phosphoproteins, 79 proteins were shared between both GLVs and systemin, 108 proteins were shared between Z3-HOL and systemin, 44 proteins were shared between Z3-HAC and systemin (Fig. 4B; Supplementary Table 2), and 112 proteins were common to both GLVs. Few proteins were dephosphoylated in response to systemin (13) and Z3-HOL (39), but Z3-HAC induced a pronounced protein dephosphorylation (255 proteins). The heatmap of P-site modifications (Fig. 4A) also shows the high number of proteins dephosphorylated in response to Z3-HAC, of which many were either phosphorylated or not significantly altered in response to Z3-HOL and systemin. A Gene Ontology (GO) analysis (Fig. 4 D,E) shows that almost all GO categories with high protein numbers for the systemin treatment also include HOL and HAC-responsive proteins (Fig. 4D), but HAC-responsive proteins were mainly dephosphorylated (Fig. 4E). Focusing in on the 79 commonly phosphorylated proteins, associated GO-categories (Supplementary Table 3) are related to signal transduction, gene expression, translation, and membrane-mediated processes.

**Fig. 4.**
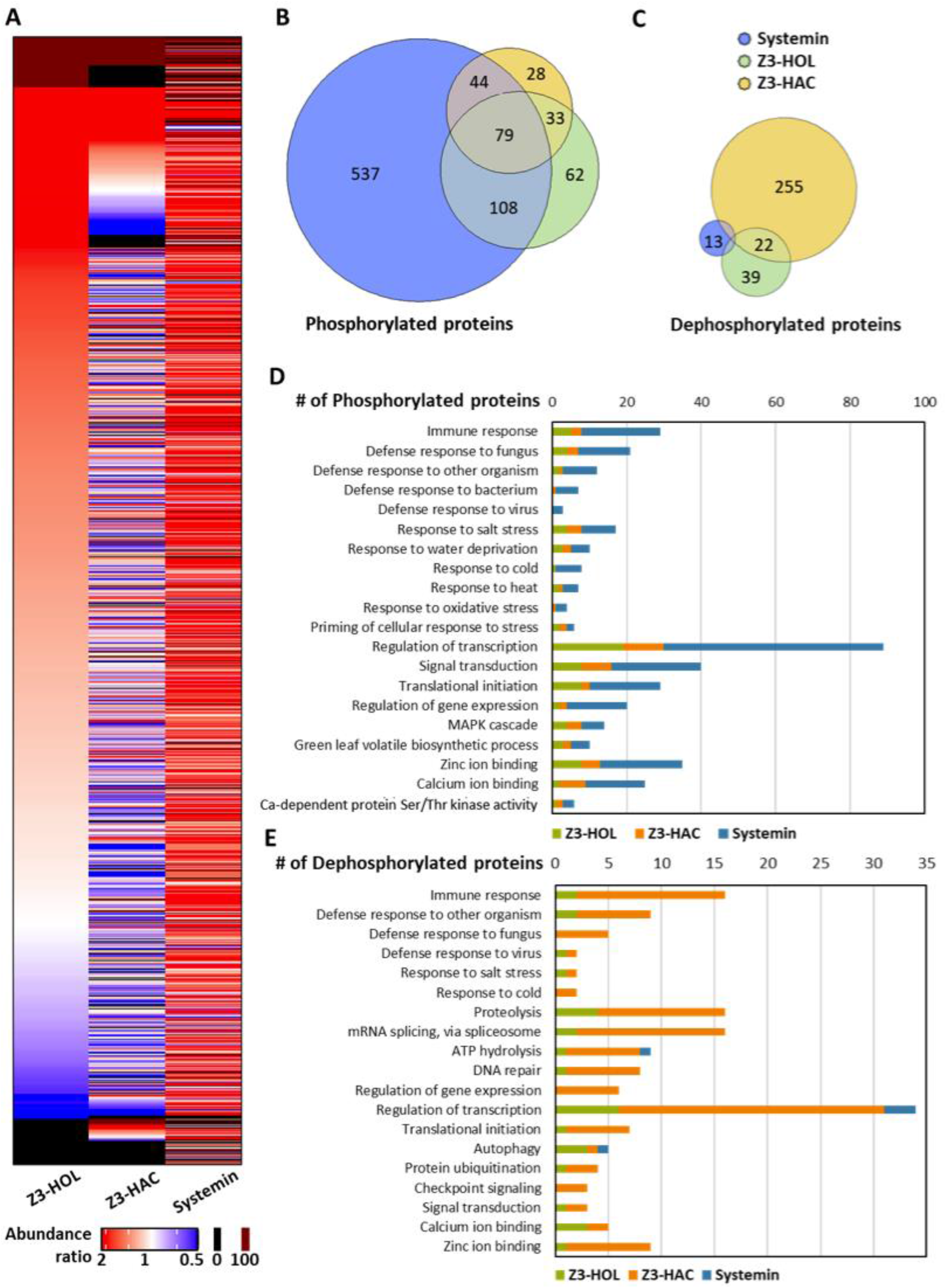
Comparison of the changes in protein phosphorylation status in response to Z3-HOL, Z3-HAC, and systemin in SP cells. **A** Heat map showing 1,502 phosphorylated and dephosphorylated proteins in response to the two GLVs and systemin. Z3-HOL-treated proteins were ranked by abundance ratio (left column). The color legend is shown at the bottom. Proteins from Z3-HAC- and systemin-treated samples were matched with Z3-HOL-treated proteins. **B, C** Venn diagrams showing overlaps of phosphorylated and dephosphorylated proteins between the three treatments, respectively. **D** Number of phosphorylated proteins and **E** number of dephosphorylated proteins in response to GLVs and systemin in selected gene ontology (GO) categories.

### Analysis of functional protein groups

The analysis of changes in the GLV-induced phosphoproteome at five minutes after treatment of SP cells revealed proteins that constitute a generic signaling pathway from perception to protein synthesis (Table 1 and Fig. 5). The same or similar proteins are well-known components of molecular pattern signaling pathways. In the following, due to the absence of an *S. peruvianum* genome and our phosphoprotein identification based on the *S. lycopersicum* reference genome, we refer to proteins from *S. peruvianum* (SP) cells with their *S. lycopersicum* taxonomic identifier (*Sl*), assuming only minor changes in sequence diversity. We identified three receptor-like kinases (*Sl*RLKs) (Solyc02g081040, Solyc09g083210, and Solyc01g102700) that were phosphorylated in response to systemin and the GLVs Z3-HAC and Z3-HOL. Two of those are PRRs characterized by a LysM domain, which is generally involved in glycan sensing. The first LysM-RLK is *Sl*LYK11 (Solyc02g081040), an ortholog of the better-known *Arabidopsis thaliana* (*At*) CERK1 (= *At*LYK1), which functions as a co-receptor for the fungal MAMP chitin. The second *Sl*LysM-RLK (Solyc09g083210) is an ortholog of Arabidopsis *At*LYK4, which engages in a chitin receptor complex consisting of *At*LYK1, *At*LYK4 and *At*LYK5 ^54^. In addition, we identified another *Sl*LysM domain protein, Solyc02g014090, which is phosphorylated in response to Z3-HOL and Z3-HAC and, based on its LysM domain, may also function in sensing of chitin or other glycans.

**Table 1.**
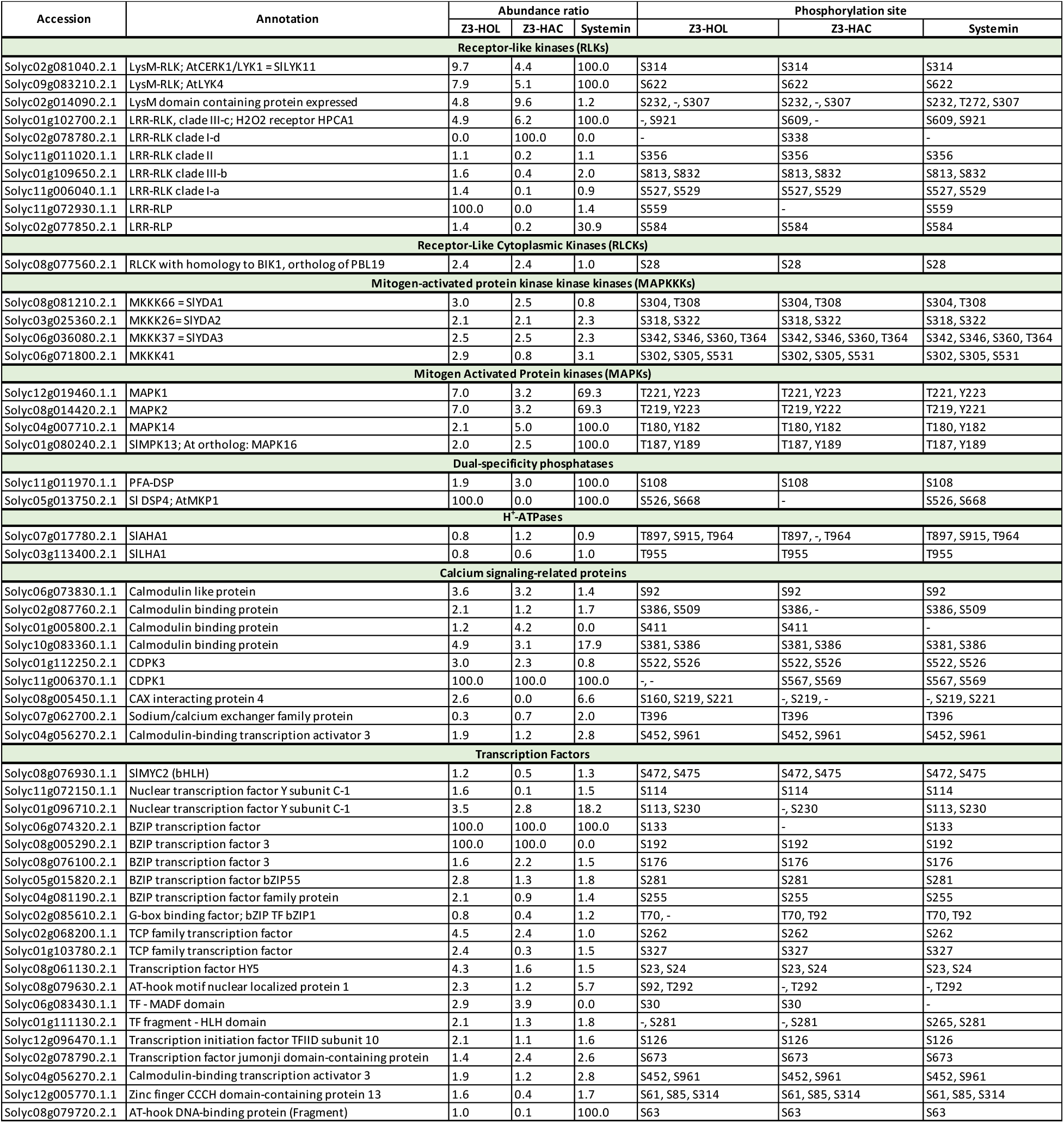
List of signaling-related proteins phosphorylated or dephosphorylated within 5 minutes in response to Z3-HOL, Z3-HAC, and systemin. Protein annotations were curated with regard to homologs in Arabidopsis.

**Fig. 5.**
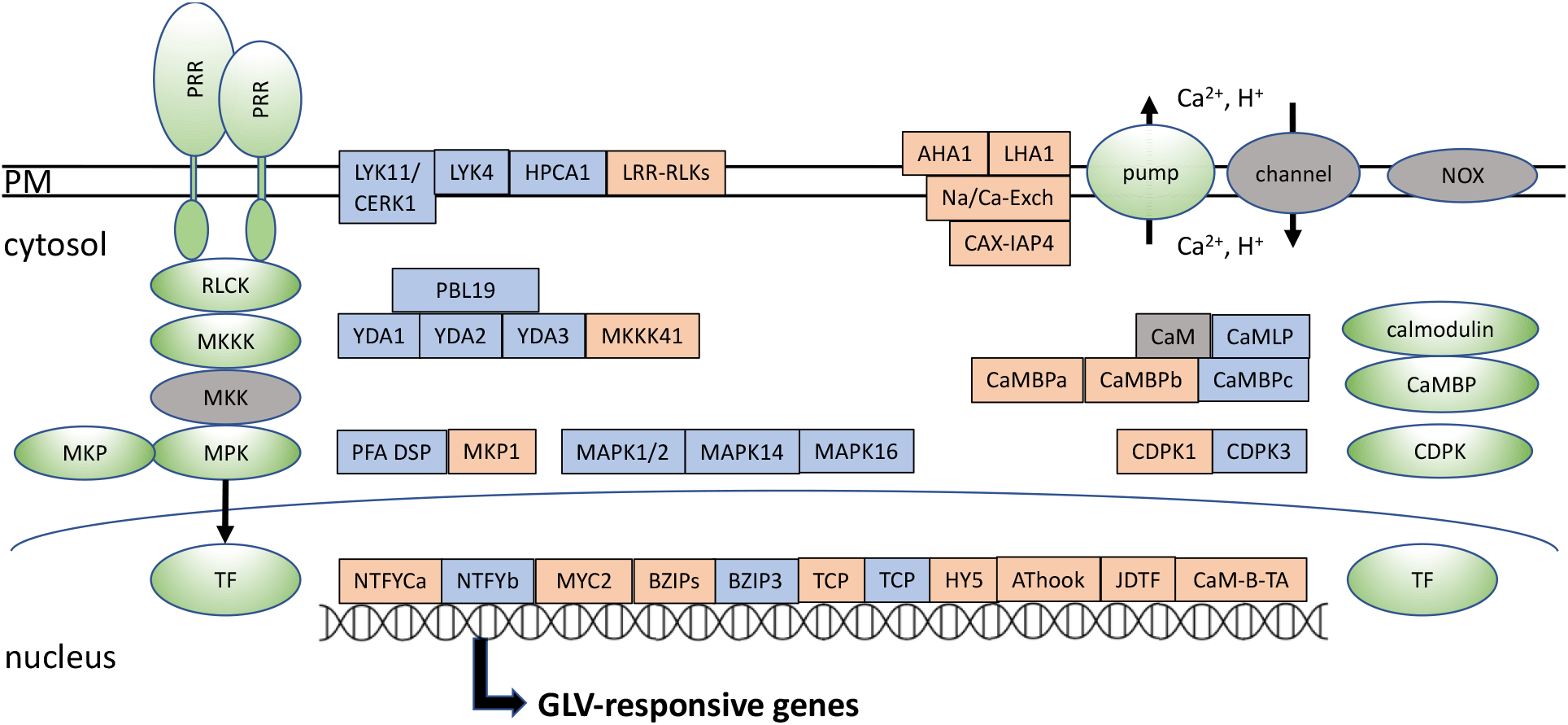
Z3-HOL- and Z3-HAC-induced changes in the phosphorylation of proteins functioning in DAMP signaling. Green ovals: Generic protein families constituting DAMP signaling pathways identified as GLV-responsive phospho-proteins. Grey ovals: Protein families not identified as phosphoproteins. Blue and orange boxes: Proteins identified as phosphoproteins responding similarly (blue) or differently (orange) to both GLVs, respectively. Acronyms for gene families not mentioned in the text (nomenclature is for *Solanum lycopersicum*): MKP– MAPK phosphatase; TF – transcription factor; NOX – NADPH oxidase; CaMBP – calmodulin binding protein; CaMBPa,b,c – small letters indicate three different proteins all featuring an IQ CaM binding domain; Na/Ca-Exch – Na^+^/Ca^2+-^exchanger; CAX-IAP4 – CAX-INTERACTING PROTEIN 4; CaMLP – CALMODULIN-LIKE PROTEIN; NTFYCa,b – two different NUCLEAR TF Y SUBUNIT C-1 proteins; TCP – TCP family TF; HY5 – ELONGATED HYPOCOTYL 5, a bZIP TF; AThook – AT-hook motif proteins; JDTF – JUMONJI DOMAIN-CONTAINING TF; CaM-B-TA – CALMODULIN-BINDING TRANSCRIPTION ACTIVATOR 3.

The third *Sl*RLK responding to both GLVs and systemin (Solyc01g102700) is an LRR-RLK and the ortholog of *At*HPCA1, the Arabidopsis receptor for H_2_O_2_. In response to H_2_O_2_, *At*HPCA1 becomes phosphorylated on three serine and three threonine residues within the intracellular domain ^19^. The phopshorylation-sites (P-sites) for the *Sl*LYK-RLKs are not conserved in Arabidopsis. The two P-sites in *Sl*HPCA1 are conserved in Arabidopsis, but were not identified when Arabidopsis plants were treated with H_2_O_2_ ^19^. Additional *Sl*RLKs or receptor-like proteins were modified differently by the two GLVs and systemin, indicating GLV-specific profiles.

RLKs, especially PRRs, directly activate receptor-like cytoplasmic kinases (RLCKs), which in turn phosphorylate and activate MAPKKKs. We found that the ser/thr kinase Solyc08g077560 is phosphorylated in response to both GLVs but not systemin. The closest homolog in Arabidopsis is PBS-LIKE 19 (PBL19) (AT5G47070), which belongs to the subfamily VII-4 of RLCKs. *At*PBL19 is known to phosphorylate MAPKKK5. Bi et al. ^55^ showed in Arabidopsis that the MAMP chitin, perceived by a CERK1-LYK5 PRR complex, activates PBL19 and other RLCK-VII4 kinases, which then activate the MAPK cascade consisting of MKKK5, MKK4/5, and MPK3/6. The Solyc08g077560 P-site at S28 is conserved in AtPBL19, but was not found in public databases.

Furthermore, we identified four GLV-responsive proteins annotated as MAPKKKs. Solyc08g081210 (*Sl*YDA1), Solyc03g025360 (*Sl*YDA2), and Solyc06g036080 (*Sl*YDA3) belong to the MEKK subfamily and are phosphorylated in response to both GLVs and systemin, except for *Sl*YDA1, which did not respond to systemin. They are orthologs of the Arabidopsis MAPKKK YODA (*At*YDA; *At*MKKK4) and close homologs of *At*MAPKKK3 and *At*MAPKKK5, which function in responses to peptide elicitors and the fungal MAMP chitin. *At*MKKK3, 4, and 5 are functionally redundant in development and immunity ^56^. Tomato *Sl*YODAs also function in immunity. The tomato *yda1* and *yda2* mutants exhibited a compromised immune response to bacterial pathogens ^57^, and all three *Sl*YDA genes are upregulated in response to *P. syringae* infection ^58^. In Arabidopsis, a target of both the YDA and MAPKKK3/5 cascades is MPK6, the ortholog of tomato MPK1 and MPK2 ^59^. The four P-sites we identified in the *Sl*YODA orthologs are conserved in AtYDA and partially in *At*MKKK5 (AT5G66850) and *At*MKKK3 (AT1G53570). The function of *Sl*MAPKKK41, which belongs to the ZIK subfamily, is unknown.

We did not identify any of the five tomato MAPKKs as phosphoproteins within our dataset. However, we found the tomato MAPKs MPK1 and MPK2 (Solyc12g019460 and Solyc08g014420) as phosphorylated on the tyrosine in their TEY activation motif in response to systemin, Z3-HOL, and Z3-HAC. This confirmed our immunoblotting results (Fig. 2), although we did not identifiy *Sl*MPK3 in our phosphoproteomics analysis. The other two GLV-responsive MAPKs, *Sl*MPK13 (Solyc01g080240) and *Sl*MPK14 (Solyc04g007710), are MAPKs with a TDY activation motif, which cannot be detected with anti-phospho-ERK (pTEpY) antibodies that we used for immunoblots (Fig. 2). While *Sl*MPK13 was phosphorylated only within the TDY phosphorylation motif, *Sl*MPK14 was also phosphorylated on serine 477 close to the C-terminus. The function of this phosphosite is unknown. *Sl*MPK1/2 and their orthologs (e.g. *At*MPK6) are the best-studied plant MAPKs and known to be involved in numerous stress responses and developmental processes ^59^, including responses to GLVs in grasses ^28,29,39^. Comparatively little is known on the functions of TDY MAPKs. Putative orthologs of *Sl*MPK13 and *Sl*MPK14 in Arabidopsis are MPKs 16 and 17, which function in ROS homeostasis ^60^.

MAPKs are dephosphorylated and inactivated by dual-specificity protein phosphatases (DSPs). We identified an *Sl*DSP (Solyc11g011970) belonging to the subfamily of PFA-DSPs. Homologs in Arabidopsis are involved in immunity ^61^ and can dephosphorylate the *Sl*MPK1/2 ortholog *At*MPK6 ^62^. Solyc11g011970 was phosphorylated in response to both GLVs and systemin. Another DSP, which is annotated as *Sl*DSP4 (Solyc05g013750), was phosphorylated in response to systemin and Z3-HOL but dephosphorylated in response to Z3-HAC. The closet homolog in Arabidopsis is *At*MKP1, a well-known MPK6-inactivating phosphatase involved in many stress responses, including those to MAMPs and pathogens ^63^. *At*MKP1 was also shown to be a negative regulator of a chitin-induced ROS burst ^64^. The two P-sites we identified in Solyc11g011970 are not conserved in Arabidopsis.

MAMPs and DAMPs, such as chitin, systemin, plant elicitor peptides (Peps), and flg22 induce an alkalinization of the extracellular pH or the growth medium of suspension-cultured cells ^43–45^, which is at least partially due to an inhibition of plasma membrane proton ATPases (H^+^-ATPases) and linked to activation of defense responses ^65^. In a phosphoproteomics analysis, Ahmad et al. ^66^ reported changes in phosphorylation of two systemin-responsive H^+^-ATPases in SP cells, Solyc03g113400 (LHA1), and Solyc07g017780 (AHA1), which correlated with a reduced H^+^-ATPase activity and alkalinization of the growth medium. This was mainly attributed to phosphorylation of the penultimate threonine residue of the two H^+^-ATPases, which stabilizes interaction with a 14-3-3 protein leading to activation of the pumps. In contrast, dephosphorylation of this P-site leads to pump inactivation. We identified the same penultimate threonine as a phosphosite in both LHA1 (T955) and AHA1 (T964). The abundance ratios for LHA1 were 0.97 fold for systemin, 0.85 for Z3-HOL, 0.63 for Z3-HAC. For AHA1, three P-sites were identified (T897, S915, T964) (Supplementary Table 4). Based on homology with the well-studied Arabidopsis H^+^-ATPases, dephosphorylation of phosphorylated T897 (pT897) results in reduced pump activity; phosphorylation of pS915 also results in reduced activity (effect of dephosphorylation unknown); and phosphorylation of pT964 (penultimate amino acid) results in activation while dephosphorylation results in inactivation ^67^. Our phosphorylation data for T897 and T964 indicate that systemin and Z3-HOL induced reduction in AHA1 activity, at least slightly, while Z3-HAC induced an increase in activity. The dephosphorylation of LHA1 (T955) (0.63 fold) in response to Z3-HAC indicates that our measured acidification response (Fig.2) is mainly due to increased activity of AHA1 or additional unidentified proton ATPases. It is also important to note that our analysis provides a snapshot at 5 min after treatment, while the actual (de-)phosphorylation of the proteome is a dynamic process ^66^.

Another common early response to stress is the activation of calcium signaling, which is initiated by Ca^2+^ ions entering the cytoplasm via channels in the plasma membrane and organellar membranes, generating a spike in the cytoplasmic Ca^2+^ concentration. Cytoplasmic Ca^2+^ ions are then recognized by calcium sensors, such as calmodulin and calcium-dependent protein kinases (CDPKs), which integrate with other signaling processes like proton fluxes and MAPK activation to specify a response. Cytosolic Ca^2+^ concentrations are reset by calcium pumps which transport Ca^2+^ ions back into storage organelles. GLV-induced modifications of P-sites were identified for a calmodulin-like protein; three proteins containing an IQ calmodulin binding domain; CDPK1 and CDPK3; a CAX-interacting protein 4 (CAX is a Ca^2+^/H^+^ -exchanger that mediates Ca^2+^ efflux); and a Na^+^/Ca^2+^-exchanger (ortholog of Arabidopsis NCX2), which functions as a pump and also mediates Ca^2+^ efflux from the cytosol. Lastly, phosphorylation of a calmodulin-binding transcriptional activator (Solyc04g056270) was changed in response to systemin and E3-HOL. Taken together, we found evidence for GLV-induced changes in the phosphorylation status of many components of calcium signaling, except for channel proteins.

Transcription factors (TFs) are targets of signaling pathways that involve MAPKs and ion fluxes. We identified GLV-induced changes in the phosphorylation status of 20 putative TFs. 12 TFs were phosphorylated in response to Z3-HOL, and 6 in response to Z3-HAC. In addition, 7 TFs were dephosphorylated in response to Z3-HAC, including the well-known TF MYC2 (Solyc08g076930), which is a master regulator of JA-responsive genes ^68^. A NUCLEAR FACTOR YC (Solyc11g072150.1.1) showed a similar response, while another NUCLEAR FACTOR YC (Solyc01g096710.2.1) was the only TF phosphorylated in response to all three treatments. Comparing all 20 TFs, each treatment induced a specific profile, which could result in GLV-specific gene expression profiles. This also shows that GLV-induced modifications of TFs happen very rapidly within five minutes.

DAMPs and MAMPs induce a profound change in the dynamics of gene expression ^69^and protein function through posttranslational modifications ^66,70^, including changes of the phosphorylation status of proteins involved in gene expression, translation, protein degradation, and primary metabolism. They are presumably involved in carrying out the transition from cellular homeostasis to stress. We identified many proteins related to these categories in the phosphoproteome of GLV-treated SP cells (Supplementary Table 4). They include proteins annotated as histones and histone-modifying enzymes, such as a histone-lysine N-methyltransferase (Solyc09g072890) for Z3-HOL and systemin, and a chromatin remodeling complex subunit (Solyc01g079700) as well as a histone deacetylase 2a-like protein (Solyc10g085560) for all three treatments. Another category contains transcription and translation-related proteins, such as mRNA processing-related proteins (e.g., for splicing and polyadenylation), ribosomal proteins, elongation factors, and chaperones. In addition, we identified phosphoproteins involved in membrane trafficking (endo- and exocytosis), such as AP1 and AP2 complex subunits, SNARE proteins, and Ras-related Rab proteins. This may include endo- and exocytosis of receptor proteins ^71^ and reflect the cellular localization dynamics underlying major changes in protein synthesis and trafficking for establishing a defense-related state. An additional set of proteins typically involved in stress responses are proteins involved in redox processes such as detoxification of reactive oxygen species. Redox-related proteins phosphorylated in response to GLVs include thioredoxin-related proteins, peroxidases, and ROS scavenger enzymes like ascorbate peroxidase, superoxide dismutase, and glutathione S-transferase. It should be noted that we neither detected an oxidative burst (Supplementary Fig. 2) nor phosphorylation of an NADPH oxidase in response to GLVs. Proteins functioning in the ubiquitin-proteasome system were also phosphorylated in response to the GLVs and systemin. Some of the proteins shown in Supplementary Table 4 are phosphorylated in response to both GLVs and systemin, but most proteins only respond to a subset of the three treatments, indicating a GLV-specific profile.

## Discussion

Taking a phosphoproteomics approach, we found that the phosphorylation state of a large number of proteins is rapidly changed in SP cells in response to the perception of the two tomato GLVs Z3-HOL and Z3-HAC, which are released in response to wounding and herbivory. Many of the rapidly modified proteins function in signal transduction from the membrane to the nucleus, including RLKs, components of MAPK and calcium signaling, and transcriptional regulators (Fig. 5). Other proteins are involved in basic cellular processes that prepare cells for a transition from an unstressed to a stressed state, such as proteins involved in transcription and translation (Supplementary Table 4). For most proteins, it is unknown what the consequences of our observed changes in phosphorylation are. In general, phosphorylation or dephosphorylation of a protein can have multiple dynamic effects, such as its ability to interact with binding partners, changing of its cellular localization, and activation or inactivation in case of an enzyme. In addition, our five-minute snapshot has to be considered within a temporal process, with any changes in phosphorylation progressing at earlier or later times.

We also demonstrated directly that GLVs induce a change in the medium pH of SP cells and MAPK phosphorylation at the TEY activation motif (Fig. 2). Both responses are typical PRR-mediated responses, and phosphorylation changes of MAPKs and H^+^-ATPases that generate the proton gradient across the plasma membrane were also identified in our phosphoproteomics data. As a reference for PRR-mediated signaling we included the DAMP systemin. An earlier phosphoproteomics study of responses to systemin in SP cells also included a five-minute time point and obtained similar results ^66^. Systemin induced changes in the phosphorylation of many more proteins than the two GLVs. Systemin also induced a stronger or different pH and MAPK response as compared to the GLVs. Since GLVs are known to induce a priming state in plants rather than a full-fledged defense response, the quantitative and qualitative differences we found between systemin and GLVs may represent these differences and may provide insights into molecular events underlying the priming phenomenon.

Notwithstanding many commonalities between Z3-HOL and Z3-HAC (112 phosphorylated proteins), we also detected differences between the two GLVs, e.g. with regard to the overlap with systemin-induced phosphorylations and the much higher number of dephosphorylated proteins in response to Z3-HAC (Fig. 4B,C). This correlates with the biphasic pH and MAPK response that is specific for Z3-HAC and has so far not been identified for other defense signals that induce a pH change (Fig. 2). Therefore, it is likely that the two GLVs co-opt PRR-mediated signaling pathways, but each GLV in a specific way, possibly resulting in different output responses. It is also noteworthy that a ROS burst, which is a common component of PRR-mediated signaling, could not be detected in response to GLVs (Supplementary Fig. 2). Accordingly, GLV treatment did not result in NADPH oxidase phosphorylation. However, systemin also did not induce an oxidative burst ^46,49^, although it induced phosphorylation of the tomato homolog of Arabidopsis RBOHD (Solyc03g117980), which was also identified by Ahmad et al. ^66^. This indicates that additional factors contribute to NADPH oxidase activity.

Since it is unknown how GLVs are perceived by plant cells, identification of RLK phosphorylation in response to GLVs may indicate a direct activation of these RLKs by GLVs. This would imply a highjacking of RLKs dedicated to specific ligands such as the MAMP chitin. GLVs induce phosphorylation of both components of the chitin-receptor complex, a CERK1 homolog (SlLYK11) and AtLYK4 homolog (Solyc09g083210). In addition, GLVs induced phosphorylation of the tomato ortholog of the Arabidopsis H_2_O_2_ receptor HPCA1. HPCA1 plays a critical role in the generation of a systemic stress-induced ROS wave, which also depends on RBOHD and Ca^2+^ signaling ^72^ and may be independent of an oxidative burst. Upregulation of RLKs in response to stress has been shown in transcriptomics studies, including in response to GLVs ^7,39,73^. In Arabidopsis, increased CERK1 protein levels were identified as part of a priming response to the salicylic acid agonist BTH ^74^ and it was proposed that increased PRR presence in the plasma membrane results in stronger and faster responses to molecular patterns in primed plants ^75^. Also, in response to flg22, oligogalacturonides, and systemin ^66,70,76^, various RLKs were identified as phosphoproteins, although the cognate receptors for the elicitors (FLS2, WAK1, and SYR1) were not. This indicates that receptor expression and posttranslational modification is a general response to many stress signals and happens in a ligand-independent manner. A possible function would be to increase sensitivity to anticipated stress signals. In contrast, after binding of flg22 to the FLS2 receptor, FLS2 is internalized and degraded, and endocytosis depends on a phosphorylation site in the FLS2 intracellular domain ^77^. RLK phosphorylation could also be the consequence of a network effect through which RLKs interact to specify and balance responses ^78^. Responses to fungi and the fungal MAMP chitin were prominent among the GLV-responsive signaling proteins. Besides the above-mentioned RLKs, we identified MPK1/2, which are also known to respond to chitin ^47^, as well as the transcription factor SlMYC2 ^79^. Many other GLV-responsive signaling proteins were identified that belong to protein categories which play a role in chitin signaling, including MAPKKKs, the tomato ortholog of the RLCK AtPBL19, DSPs, proton ATPases, calcium signaling-related proteins, and transcription factors (Table 1). A metanalysis showed that attack by fungi has a stronger effect on GLV release by plants than wounding or herbivory ^1^. Therefore, it may be adaptive for the plant to interpret emitted GLVs as a sign of fungal attack. Since fungal attacks often start locally, this could have evolved as an airborne intraplant communication system.

While much has been learned about GLV emissions and their role in intra- and interplant communication, very little is known about signaling and perception. Our data provide novel insights into GLV signaling suggesting that GLVs co-opt PRR-mediated signaling pathways, and that the GLV-induced phosphoproteome dynamics may represent an early signaling stage leading to priming.

## Methods

### Chemicals and Stock Solutions

The stock solutions for the GLVs (*Z*)-3-hexen-1-ol (Z3-HOL, 98%; Acros Organics) and (*Z*)-3-hexenyl acetate (Z3-HAC, 99%; Alfa Aesar) were solved in ethanol (100%, Decon Labs, Inc.) for pH, MAPK activity, and phosphoproteomic experiments and dimethyl sulfoxide (DMSO, Fisher Scientific) for oxidative burst experiments. The stock solutions of peptide elicitors systemin and flg22 (GenScript) were solved in water.

### Suspension-cultured Cells and Plants

*Solanum peruvianum* suspension-cultured cells (SP cells) were originally generated by L. Nover in 1982 ^80^ and established for testing MAMP responses by the Boller group ^81^. SP cells were grown in 125-mL Erlenmeyer flasks on an orbital shaker (175 rpm) under ambient conditions. Cells were subcultured every 12 days and used for experiments 10 days after subculturing. Tomato plants (*S. lycopersicum* var. *Microtom*) were grown in AR66L growth chambers (Percival Scientific) on a 16 h light (130 ± 20 μE m^−2^ s^−1^; 27°C) and 8 h dark (22°C) in Jiffy peat pellets. Two- to three-week-old plants were used for oxidative burst experiments.

### pH Assay

Medium pH was measured as described ^45^. Briefly, SP cells (1.5 mL) were transferred into wells of non-treated, sterilized 12-well tissue culture plates (Avantor) and shaken (175 rpm) on an orbital shaker under ambient conditions. After a 1 h adjustment period, the cells were treated with a final concentration of 10 nM systemin, 0.7% ethanol, 5 – 28.5 mM Z3-HOL, or 270 μM – 4.2 mM Z3-HAC. After treatment, the medium pH was monitored using a pH probe (Mettler Toledo). To prevent cross-contamination by volatilized GLVs, Z3-HOL, Z3-HAC, and control treatments were each performed on separate plates.

### MAPK Phosphorylation Assay

MAPK assays were performed as described (Hann et al., 2014). Briefly, proteins from SP cells treated with a final concentration of 3 nM systemin, 1% ethanol, 28.5 mM Z3-HOL, or 4.2 mM Z3-HAC were extracted as described (Holley et al., 2003). 20 μg of protein (15-well gels; Bio-Rad Mini-PROTEAN tetra system) was analyzed for MAPK phosphorylation by immunoblotting using the primary antibody anti-pERK MAPK (Phospho-p44/p42 MAPK, ERK1/2,Thr202/Tyr204, D13.14.4E; Cell Signaling Technology, Danvers, MA, USA) at a dilution of 1:2,500 and the secondary antibody (Goat anti-rabbit IgG (H+L)-HRP conjugate; Bio-Rad) at a 1:20,000 dilution, followed by a chemiluminescence assay with the Clarity™ Western ECL substrate (Bio-Rad). Membranes were stained with Coomassie brilliant blue to ensure equal protein transfer and loading.

### Oxidative Burst Assay

Tomato leaf disks were cut using a 4-mm cork borer, and one disk was placed adaxial side up into the wells of a 96-well plate (Lumitrac 200; Greiner Bio-One) containing 200 μL of water. After an incubation in the dark under ambient temperature for 12- 16 hours, the water was removed and replaced with 100 μL of an aqueous solution containing 7.8 μg/mL L-012 (Sigma Aldrich) and 20 μg/mL horseradish peroxidase (Alfa Aesar) each, plus 10 nM flg22 (solved in DMSO), 10 nM systemin (solved in DMSO), 0.6% DMSO (blank), 500 μM – 28.5 mM Z3-HOL, or 270 μM – 4.2 mM Z3-HAC. Chemiluminescence was measured using a Synergy HTX Multi-Mode Microplate Reader (BioTek) as described (Hann et al., 2014).

### Cell treatment, cell lysis, and protein digestion for phosphoproteomic analysis

SP cells were treated with 15 μL of Z3-HOL or Z3-HAC, solved in ethanol, at a final concentration of 28.5 mM and 4.2 mM, respectively. Ethanol (15 μL) was added to the control group, and a treatment with 10 nM systemin served as a reference. Treated SP cells were lysed using the method described by Mullis et al.^51^. Briefly, 1 mL of RIPA buffer (Thermo Scientific Pierce, Rockford, IL) containing 1× Halt Protease and Phosphatase Inhibitor Cocktail (Thermo Scientific Pierce) was added to each sample. Proteins were reduced with 10 mM TCEP at 56 °C for 30 min and then alkylated with 40 mM iodoacetamide for 30 min in the dark at room temperature. Ice-cold acetone was used to precipitate the proteins at −80 °C and −20 °C, respectively. Then, the samples were centrifuged, and protein pellets were collected. The pellets were resuspended in 50 mM ammonium bicarbonate, followed by trypsin digestion using MS Grade Trypsin Protease (Thermo Scientific Pierce). Afterward, the samples were acidified and centrifuged. Supernatants were collected and stored at –80 °C.

### Phosphopeptide enrichment of SP protein digests

Phosphopeptides in the protein digests were enriched using the automated robotic platform-based method described by Mullis et al. ^51^. Protein digests were thawed on ice and 20 μL aliquots were diluted to the final volume of 200 μL with 1% TFA before they were transferred to a 96-well plate. Then, samples were desalted using 5 mg RP 300 μL IMCStips (IMCS, Irmo, SC), followed by phosphopeptide enrichment using 10 μg PolyTi 300 μL IMCStips (IMCS, Irmo, SC). The automated peptide desalting and phosphopeptide enrichment was performed on a Hamilton STAR system following the previously published workflow ^51^. The eluted solutions were lyophilized and stored at −80 °C until further analysis.

### Mass Spectrometry (MS) Analysis

Enriched phosphopeptide samples were redissolved in 4.8 μL of formic acid and acetonitrile mixture, 0.25% and 3% (v/v), respectively. A Dionex Ultimate 3000 nano-flow liquid chromatograph was used. 4 μL of sample was first injected into a cartridge trap column (Thermo Fisher Scientific), then transferred to a capillary analytical column for separation (PepMap RSLC C18, EasySpray format, 15 cm by 75 μm packed with 3 um 100Å particles). The EasySpray nano-spray source was interfaced to a Thermo Q-Exactive HF-X quadrupole-orbitrap mass spectrometer which was operated using a standard Top-6 data dependent acquisition (DDA) protocol at Nevada Proteomics Center (Reno, NV).

The raw data were searched against a *Solanum lycopersicum* protein database (ITAG 2.3) using SequestHT algorithms on Proteome Discoverer 2.2 (Thermo Fisher). The peptide precursor mass tolerance was set to 10 ppm, and the fragment mass tolerance was set to 0.6 Da. Search criteria included a static modification of cysteine residues of +57.0214 Da and a variable modification of +15.995 Da to include potential oxidation of methionine and a modification of +79.966 Da on serine, threonine, or tyrosine for the identification of phosphorylation (three modifications allowed per peptide). Searches were performed with full tryptic digestion and allowed a maximum of two missed cleavages on the peptides analyzed from the sequence database. False discovery rates (FDRs) were set to less than 1% for each analysis at the peptide-spectrum match (PSM) level. Phosphorylation site localization from collision-induced decomposition (CID) spectra was determined by PhosphoRS on Proteome Discoverer 2.2. Heatmap and proportional Venn diagrams were generated using R. Abundance ratios were determined by using the ethanol-treated group as a control for GLVs, and the untreated group as a control for systemin. An abundance ratio of 100 was defined as an abundance value present only in the treatment, but not in the corresponding control. An abundance ratio of zero was defined as an abundance value only found in the control group, but not in the treatment.

## Data availability

All relevant data supporting our findings are available within Supplementary Material for this work.

## Acknowledgements

This work was supported by a grant from the National Science Foundation to Johannes Stratmann and Qian Wang (NSF-IOS 2051699), and by an ASPIRE I grant from the Vice President for Research at the University of South Carolina.

## Author contributions

S.T. and K.F. performed most of the experiments and analyzed the data. B.T.M. performed intial phosphoproteomics analysis. J.R. performed immunoblots. H.N. helped with annotation of phosphoproteins. Q.W. and J.W.S. conceived, designed, and directed the project and wrote the manuscript.

## Supplementary Material

- **Supplementary Figure 1**. Early and late responses of SP cells to GLVs and fusicoccin

- **Supplementary Figure 2**. GLVs do not induce an oxidative burst in leaf discs of tomato plants.

- **Supplementary Table 1**. Abundance ratios for all proteins phosphorylated and dephosphorylated in response to Z3-HOL, Z3-HAC, and systemin

- **Supplementary Table 2**. Detailed information for phosphoproteins in all 14 sectors of the Venn diagram shown in Fig. 4

- **Supplementary Table 3**. GO-categories for 79 proteins phosphorylated in response to systemin, Z3-HOL, and Z3-HAC

- **Supplementary Table 4**. AHA1 phosphorylation sites and abundance ratios

- **Supplementary Table 5**. List of proteins phosphorylated or dephosphorylated within 5 minutes in response to Z3-HOL, Z3-HAC, and systemin.

## References

1 Ameye, M. et al. Green leaf volatile production by plants: a meta-analysis. New Phytol. 220, 666–683, doi:10.1111/nph.14671 (2018).

2 Degenhardt, D. C., Refi-Hind, S., Stratmann, J. W. & Lincoln, D. E. Systemin and jasmonic acid regulate constitutive and herbivore-induced systemic volatile emissions in tomato, Solanum lycopersicum. Phytochemistry 71, 2024–2037 (2010).

3 Joo, Y. et al. Herbivory elicits changes in green leaf volatile production via jasmonate signaling and the circadian clock. Plant, Cell Environ 42, 972–982, doi:10.1111/pce.13474 (2019).

4 Heil, M. & Land, W. G. Danger signals – damaged-self recognition across the tree of life. Front. Plant Sci. 5, doi:10.3389/fpls.2014.00578 (2014).

5 Heil, M. Herbivore-induced plant volatiles: targets, perception and unanswered questions. New Phytol. 204, 297–306, doi:https://doi.org/10.1111/nph.12977 (2014).

6 Meents, A. K. & Mithofer, A. Plant-Plant Communication: Is there a role for volatile damage-associated molecular patterns? Front. Plant Sci. 11, doi:https://doi.org/10.3389/fpls.2020.583275 (2020).

7 Yamauchi, Y., Matsuda, A., Matsuura, N., Mizutani, M. & Sugimoto, Y. Transcriptome analysis of Arabidopsis thaliana treated with green leaf volatiles: possible role of green leaf volatiles as self-made damage-associated molecular patterns. J. Pestic. Sci. 43, 207–213, doi:10.1584/jpestics.D18-020 (2018).

8 Heil, M. & Silva Bueno, J. C. Within-plant signaling by volatiles leads to induction and priming of an indirect plant defense in nature. Proc. Natl. Acad. Sci. U. S. A. 104, 5467–5472, doi:10.1073/pnas.0610266104 (2007).

9 Karban, R., Shiojiri, K., Huntzinger, M. & McCall, A. C. Damage-induced resistance in sagebrush: volatiles are key to intra- and interplant communication. Ecology 87, 922–930, doi:10.1890/0012-9658(2006)87[922:drisva]2.0.co;2 (2006).

10 Kim, J. & Felton, G. W. Priming of antiherbivore defensive responses in plants. Insect Sci. 20, 273–285, doi:10.1111/j.1744-7917.2012.01584.x (2013).

11 Engelberth, J., Alborn, H. T., Schmelz, E. A. & Tumlinson, J. H. Airborne signals prime plants against insect herbivore attack. Proc. Natl. Acad. Sci. U. S. A. 101, 1781–1785 (2004).

12 Matsui, K. & Engelberth, J. Green Leaf Volatiles—The forefront of plant responses against biotic attack. Plant Cell Physiol., doi:10.1093/pcp/pcac117 (2022).

13 Ranf, S. Sensing of molecular patterns through cell surface immune receptors. Curr. Opin. Plant Biol. 38, 68–77, doi:10.1016/j.pbi.2017.04.011 (2017).

14 Sato, K. et al. Insect olfactory receptors are heteromeric ligand-gated ion channels. Nature 452, 1002–1006, doi:10.1038/nature06850 (2008).

15 Moriyama, E. N., Strope, P. K., Opiyo, S. O., Chen, Z. & Jones, A. M. Mining the *Arabidopsis thaliana* genome for highly-divergent seven transmembrane receptors. Genome Biol. 7, R96, doi:10.1186/gb-2006-7-10-r96 (2006).

16 Adebesin, F. et al. Emission of volatile organic compounds from petunia flowers is facilitated by an ABC transporter. Science 356, 1386–1388, doi:10.1126/science.aan0826 (2017).

17 Widhalm, J. R., Jaini, R., Morgan, J. A. & Dudareva, N. Rethinking how volatiles are released from plant cells. Trends Plant Sci. 20, 545–550, doi:10.1016/j.tplants.2015.06.009 (2015).

18 Wang, Z.-Y., Seto, H., Fujioka, S., Yoshida, S. & Chory, J. BRI1 is a critical component of a plasma-membrane receptor for plant steroids. Nature 410, 380–383, doi:10.1038/35066597 (2001).

19 Wu, F. et al. Hydrogen peroxide sensor HPCA1 is an LRR receptor kinase in Arabidopsis. Nature 578, 577–581, doi:10.1038/s41586-020-2032-3 (2020).

20 Kutschera, A. & Ranf, S. The multifaceted functions of lipopolysaccharide in plant-bacteria interactions. Biochimie 159, 93–98, doi:10.1016/j.biochi.2018.07.028 (2019).

21 Kato, H. et al. Pathogen-derived 9-methyl sphingoid base is perceived by a lectin receptor kinase in Arabidopsis. bioRxiv, 2021.2010.2018.464766, doi:10.1101/2021.10.18.464766 (2021).

22 Hall, B. P., Shakeel, S. N. & Schaller, G. E. Ethylene Receptors: ethylene perception and signal transduction. J. Plant Growth Regul. 26, 118–130, doi:10.1007/s00344-007-9000-0 (2007).

23 Heil, M., Lion, U. & Boland, W. Defense-inducing volatiles: in search of the active motif. J. Chem. Ecol. 34, 601–604, doi:10.1007/s10886-008-9464-9 (2008).

24 Zebelo, S. A., Matsui, K., Ozawa, R. & Maffei, M. E. Plasma membrane potential depolarization and cytosolic calcium flux are early events involved in tomato (Solanum lycopersicon) plant-to-plant communication. Plant Sci. 196, 93–100, doi:10.1016/j.plantsci.2012.08.006 (2012).

25 Matsui, K. A portion of plant airborne communication is endorsed by uptake and metabolism of volatile organic compounds. Curr. Opin. Plant Biol. 32, 24–30, doi:10.1016/j.pbi.2016.05.005 (2016).

26 Asai, N., Nishioka, T., Takabayashi, J. & Furuichi, T. Plant volatiles regulate the activities of Ca2+ -permeable channels and promote cytoplasmic calcium transients in Arabidopsis leaf cells. Plant Signal. Behav. 4, 294–300, doi:10.4161/psb.4.4.8275 (2009).

27 Gilroy, S. et al. A tidal wave of signals: calcium and ROS at the forefront of rapid systemic signaling. Trends Plant Sci. 19, 623–630, doi:10.1016/j.tplants.2014.06.013 (2014).

28 Dombrowski, J. E. & Martin, R. C. Green leaf volatiles, fire and nonanoic acid activate MAPkinases in the model grass species *Lolium temulentum*. BMC Res. Notes 7, 807, doi:10.1186/1756-0500-7-807 (2014).

29 Dombrowski, J. E. & Martin, R. C. Activation of MAP kinases by green leaf volatiles in grasses. BMC Res. Notes 11, 79, doi:10.1186/s13104-017-3076-9 (2018).

30 Seyfferth, C. & Tsuda, K. Salicylic acid signal transduction: the initiation of biosynthesis, perception and transcriptional reprogramming. Front. Plant Sci. 5, doi:10.3389/fpls.2014.00697 (2014).

31 Herrera-Vásquez, A., Salinas, P. & Holuigue, L. Salicylic acid and reactive oxygen species interplay in the transcriptional control of defense genes expression. Front. Plant Sci. 6, doi:10.3389/fpls.2015.00171 (2015).

32 Kandoth, P. K. et al. Tomato MAPKs LeMPK1, LeMPK2, and LeMPK3 function in the systemin-mediated defense response against herbivorous insects. Proc. Natl. Acad. Sci. U. S. A. 104, 12205–12210, doi:10.1073/pnas.0700344104 (2007).

33 Erb, M. & Reymond, P. Molecular Interactions Between Plants and Insect Herbivores. Annu. Rev. Plant Biol. 70, 527–557, doi:10.1146/annurev-arplant-050718-095910 (2019).

34 Engelberth, J., Seidl-Adams, I., Schultz, J. C. & Tumlinson, J. H. Insect elicitors and exposure to green leafy volatiles differentially upregulate major octadecanoids and transcripts of 12-oxo phytodienoic acid reductases in Zea mays. Mol. Plant Microbe Interact. 20, 707–716, doi:10.1094/MPMI-20-6-0707 (2007).

35 Paudel Timilsena, B., Seidl-Adams, I. & Tumlinson, J. H. Herbivore-specific plant volatiles prime neighboring plants for nonspecific defense responses. Plant, Cell Environ., doi:10.1111/pce.13688 (2019).

36 Su, Q. et al. Defence priming in tomato by the green leaf volatile (Z)-3-hexenol reduces whitefly transmission of a plant virus. Plant, Cell Environ. 43, 2797–2811, doi:10.1111/pce.13885 (2020).

37 Engelberth, J., Contreras, C. F., Dalvi, C., Li, T. & Engelberth, M. Early transcriptome analyses of Z-3-Hexenol-treated *Zea mays* revealed distinct transcriptional networks and anti-herbivore defense potential of green leaf volatiles. PLoS One 8, e77465, doi:10.1371/journal.pone.0077465 (2013).

38 Kishimoto, K., Matsui, K., Ozawa, R. & Takabayashi, J. Volatile C6-aldehydes and Allo-ocimene Activate Defense Genes and Induce Resistance against *Botrytis cinerea* in *Arabidopsis thaliana*. Plant Cell Physiol. 46, 1093–1102, doi:10.1093/pcp/pci122 (2005).

39 Dombrowski, J. E. et al. Transcriptome analysis of the model grass Lolium temulentum exposed to green leaf volatiles. BMC Plant Biol. 19:222 https://doi.org/10.1186/s12870-019-1799-6 (2019)

40 Mirabella, R. et al. The Arabidopsis her1 mutant implicates GABA in E-2-hexenal responsiveness. Plant J. 53, 197–213, doi:10.1111/j.1365-313X.2007.03323.x (2008).

41 Scala, A. et al. Forward genetic screens identify a role for the mitochondrial HER2 in E-2-hexenal responsiveness. Plant Mol. Biol. 95, 399–409, doi:10.1007/s11103-017-0659-8 (2017).

42 Tarkowski, L. P., Signorelli, S. & Hofte, M. gamma-Aminobutyric acid and related amino acids in plant immune responses: Emerging mechanisms of action. Plant Cell Environ 43, 1103–1116, doi:10.1111/pce.13734 (2020).

43 Felix, G., Duran, J. D., Volko, S. & Boller, T. Plants have a sensitive perception system for the most conserved domain of bacterial flagellin. Plant J. 18, 265–276 (1999).

44 Huffaker, A., Pearce, G. & Ryan, C. A. An endogenous peptide signal in Arabidopsis activates components of the innate immune response. Proc. Natl. Acad. Sci. U. S. A. 103, 10098–10103 (2006).

45 Stratmann, J., Scheer, J. & Ryan, C. A. Suramin inhibits initiation of defense signaling by systemin, chitosan and pmg-elicitor in suspension cultured *Lycopersicon peruvianum* cells. Proc. Natl. Acad. Sci. USA 97, 8862–8867, doi:https://doi.org/10.1073/pnas.97.16.8862 (2000).

46 Hann, C. T., Bequette, C. J., Dombrowski, J. E. & Stratmann, J. W. Methanol and ethanol modulate responses to danger- and microbe-associated molecular patterns. Front. Plant Sci. 5, doi:10.3389/fpls.2014.00550 (2014).

47 Holley, S. R., Yalamanchili, R. D., Moura, S. D., Ryan, C. A. & Stratmann, J. W. Convergence of signaling pathways induced by systemin, oligosaccharide elicitors, and ultraviolet-B radiation at the level of mitogen-activated protein kinases in *Lycopersicon peruvianum* suspension-cultured cells. Plant Physiol. 132, 1728–1738 (2003).

48 Rahman, H., Wang, X. Y., Xu, Y. P., He, Y. H. & Cai, X. Z. Characterization of tomato protein kinases embedding guanylate cyclase catalytic center motif. Sci Rep-Uk 10, https://doi.org/10.1038/s41598-020-61000-7 (2020).

49 Xu, S. et al. Tomato PEPR1 ORTHOLOG RECEPTOR-LIKE KINASE1 Regulates Responses to Systemin, Necrotrophic Fungi, and Insect Herbivory. Plant Cell 30, 2214–2229, doi:10.1105/tpc.17.00908 (2018).

50 Wu, X., Sklodowski, K., Encke, B. & Schulze, W. X. A kinase-phosphatase signaling module with BSK8 and BSL2 involved in regulation of sucrose-phosphate synthase. J Proteome Res. 13, 3397–3409, doi:10.1021/pr5003164 (2014).

51 Mullis, B. T. et al. Automating Complex, Multistep Processes on a Single Robotic Platform to Generate Reproducible Phosphoproteomic Data. SLAS DISCOVERY: Advancing the Science of Drug Discovery 25, 277–286, doi:10.1177/2472555219878152 (2020).

52 Hsu, C. C., Arrington, J. V. & Tao, W. A. Universal Sample Preparation Workflow for Plant Phosphoproteomic Profiling. Methods Mol. Biol. 2358, 93–103, doi:10.1007/978-1-0716-1625-3_6 (2021).

53 Iliuk, A. B., Martin, V. A., Alicie, B. M., Geahlen, R. L. & Tao, W. A. In-depth analyses of kinase-dependent tyrosine phosphoproteomes based on metal ion-functionalized soluble nanopolymers. Mol. Cell Proteomics 9, 2162–2172, doi:10.1074/mcp.M110.000091 (2010).

54 Xue, D. X., Li, C. L., Xie, Z. P. & Staehelin, C. LYK4 is a component of a tripartite chitin receptor complex in *Arabidopsis thaliana*. J. Exp. Bot. 70, 5507–5516, doi:10.1093/jxb/erz313 (2019).

55 Bi, G. Z. et al. Receptor-Like Cytoplasmic Kinases Directly Link Diverse Pattern Recognition Receptors to the Activation of Mitogen-Activated Protein Kinase Cascades in Arabidopsis. Plant Cell 30, 1543–1561, doi:10.1105/tpc.17.00981 (2018).

56 Wang, Y. P., Wu, Y. Y., Zhang, H. L., Wang, P. X. & Xia, Y. J. Arabidopsis MAPKK kinases YODA, MAPKKK3, and MAPKKK5 are functionally redundant in development and immunity. Plant Physiol. 190, 206–210, doi:10.1093/plphys/kiac270 (2022).

57 Tellez, J. et al. YODA Kinase Controls a Novel Immune Pathway of Tomato Conferring Enhanced Disease Resistance to the Bacterium *Pseudomonas syringae*. Front. Plant Sci. 11, https://doi.org/10.3389/fpls.2020.584471 (2020).

58 Wu, J. et al. Genome-wide identification of MAPKK and MAPKKK gene families in tomato and transcriptional profiling analysis during development and stress response. PLoS One 9, e103032, doi:10.1371/journal.pone.0103032 (2014).

59 Zhang, M. M. & Zhang, S. Q. Mitogen-activated protein kinase cascades in plant signaling. J. Integr. Plant Biol. 64, 301–341, doi:10.1111/jipb.13215 (2022).

60 Takahashi, F., Mizoguchi, T., Yoshida, R., Ichimura, K. & Shinozaki, K. Calmodulin-Dependent Activation of MAP Kinase for ROS Homeostasis in Arabidopsis. Mol. Cell 41, 649–660, doi:10.1016/j.molcel.2011.02.029 (2011).

61 He, H. J. et al. Two Homologous Putative Protein Tyrosine Phosphatases, OsPFA-DSP2 and AtPFA-DSP4, Negatively Regulate the Pathogen Response in Transgenic Plants. Plos One 7, https://doi.org/10.1371/journal.pone.0034995 (2012).

62 Xin, J. et al. AtPFA-DSP3, an atypical dual-specificity protein tyrosine phosphatase, affects salt stress response by modulating MPK3 and MPK6 activity. Plant, Cell Environ. 44, 1534–1548, doi:10.1111/pce.14002 (2021).

63 Jiang, L. Y., Chen, Y. H., Luo, L. J. & Peck, S. C. Central Roles and Regulatory Mechanisms of Dual-Specificity MAPK Phosphatases in Developmental and Stress Signaling. Front. Plant Sci. 9, https://doi.org/10.3389/fpls.2018.01697 (2018).

64 Escudero, V. et al. Mitogen-Activated Protein Kinase Phosphatase 1 (MKP1) Negatively Regulates the Production of Reactive Oxygen Species During Arabidopsis Immune Responses. Mol. Plant-Microbe Interact. 32, 464–478, doi:10.1094/Mpmi-08-18-0217-Fi (2019).

65 Schaller, A. & Oecking, C. Modulation of plasma membrane H^+^-ATPase activity differentially activates wound and pathogen defense responses in tomato plants. Plant Cell 11, 263–272 (1999).

66 Ahmad, F. H., Wu, X. N., Stintzi, A., Schaller, A. & Schulze, W. X. The Systemin Signaling Cascade As Derived from Time Course Analyses of the Systemin-responsive Phosphoproteome. Mol. Cell. Proteomics 18, 1526–1542, doi:10.1074/mcp.RA119.001367 (2019).

67 Falhof, J., Pedersen, J. T., Fuglsang, A. T. & Palmgren, M. Plasma Membrane H(+)-ATPase Regulation in the Center of Plant Physiology. Mol. Plant 9, 323–337, doi:10.1016/j.molp.2015.11.002 (2016).

68 Du, M. M. et al. MYC2 Orchestrates a Hierarchical Transcriptional Cascade That Regulates Jasmonate-Mediated Plant Immunity in Tomato. Plant Cell 29, 1883–1906, doi:10.1105/tpc.16.00953 (2017).

69 Winkelmuller, T. M. et al. Gene expression evolution in pattern-triggered immunity within Arabidopsis thaliana and across Brassicaceae species. Plant Cell 33, 1863–1887, doi:10.1093/plcell/koab073 (2021).

70 Benschop, J. J. et al. Quantitative phosphoproteomics of early elicitor signaling in Arabidopsis. Mol. Cell. Proteomics 6, 1198–1214, doi:10.1074/mcp.M600429-MCP200 (2007).

71 Gu, Y. N., Zavaliev, R. & Dong, X. N. Membrane Trafficking in Plant Immunity. Mol. Plant 10, 1026–1034, doi:10.1016/j.molp.2017.07.001 (2017).

72 Fichman, Y., Zandalinas, S. I., Peck, S., Luan, S. & Mittler, R. HPCA1 is required for systemic reactive oxygen species and calcium cell-to-cell signaling and plant acclimation to stress. Plant Cell, doi:10.1093/plcell/koac241 (2022).

73 Mirabella, R. et al. WRKY40 and WRKY6 act downstream of the green leaf volatile E-2-hexenal in Arabidopsis. Plant J. 83, 1082–1096, doi:10.1111/tpj.12953 (2015).

74 Tateda, C. et al. Salicylic Acid Regulates Arabidopsis Microbial Pattern Receptor Kinase Levels and Signaling. Plant Cell 26, 4171–4187, doi:10.1105/tpc.114.131938 (2014).

75 Conrath, U., Beckers, G. J. M., Langenbach, C. J. G. & Jaskiewicz, M. R. Priming for Enhanced Defense. Annu. Rev. Phytopathol., Vol 53 53, 97–119, doi:10.1146/annurev-phyto-080614-120132 (2015).

76 Mattei, B., Spinelli, F., Pontiggia, D. & Lorenzo De, G. Comprehensive Analysis of the Membrane Phosphoproteome Regulated by Oligogalacturonides in *Arabidopsis thaliana*. Front. Plant Sci. 7, https://doi.org/10.3389/fpls.2016.01107 (2016).

77 Robatzek, S., Chinchilla, D. & Boller, T. Ligand-induced endocytosis of the pattern recognition receptor FLS2 in Arabidopsis. Genes Dev. 20, 537–542, doi:10.1101/gad.366506 (2006).

78 Smakowska-Luzan, E. et al. An extracellular network of Arabidopsis leucine-rich repeat receptor kinases. Nature 553, 342-+, doi:10.1038/nature25184 (2018).

79 Libault, M., Wan, J. R., Czechowski, T., Udvardi, M. & Stacey, G. Identification of 118 Arabidopsis transcription factor and 30 ubiquitin-ligase genes responding to chitin, a plant-defense elicitor. Mol. Plant-Microbe Interact. 20, 900–911, doi:10.1094/Mpmi-20-8-0900 (2007).

80 Nover, L., Kranz, E. & Scharf, K. D. Growth-Cycle of Suspension-Cultures of Lycopersicon-Esculentum and *Lycopersicon-Peruvianum*. Biochem. Physiol. Pfl. 177, 483–499, doi:Doi 10.1016/S0015-3796(82)80041-3 (1982).

81 Felix, G. & Boller, T. Systemin induces rapid ion fluxes and ethylene biosynthesis in *Lycopersicon peruvianum* cells. Plant J. 7, 381–389 (1995).

