## Supplemental Figure 1 for "Green leaf volatiles co-opt proteins involved in molecular pattern signaling in plant cells"

### Supplementary Figure 1

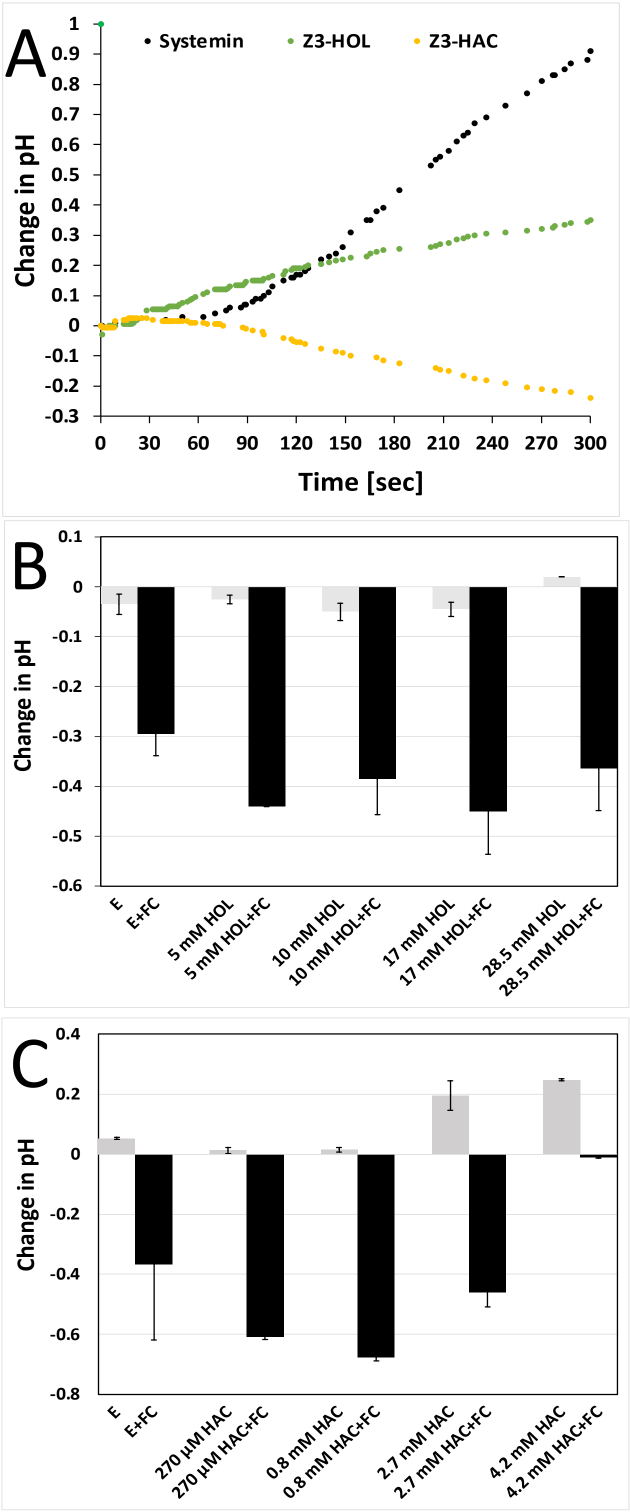


**Supplementary Figure 1. Early and late responses of SP cells to GLVs and fusicoccin. A** Systemin- and GLV-induced medium pH changes were measured every two seconds for the first five minutes (Z3-HOL = 28.5 mM, Z3-HAC = 4.2 mM, Systemin = 10 nM systemin). A representative experiment of 3 independent experiments (n = 3) is shown. **B, C** SP cells were treated with different concentrations of Z3-HOL (HOL) (**B**) or Z3-HAC (HAC) (**C**) as indicated, resulting in an increase or decrease in medium pH as shown in Fig. 1A,B. 90 min after GLV or ethanol treatment, cells were treated with 2.5 µM fusicoccin (FC). FC is a plasma membrane proton ATPase agonist that causes medium acidification. 15 min after FC treatment, the medium pH was measured. Bars represent the average and SD of two independent experiments (n = 2). Since proton ATPase activity can still be regulated 90 min after GLV treatment, the SP cells are not irreversibly damaged and can maintain a proton gradient across the plasma membrane. Note that 30 min after HAC treatment, the medium pH is increasing as shown in Fig. 1B, which is reflected in the increased pH induced by 2.7 and 4.2 mM HAC in panel C.
