## Supplemental Figure 2 for "Green leaf volatiles co-opt proteins involved in molecular pattern signaling in plant cells"

### Supplementary Figure 2

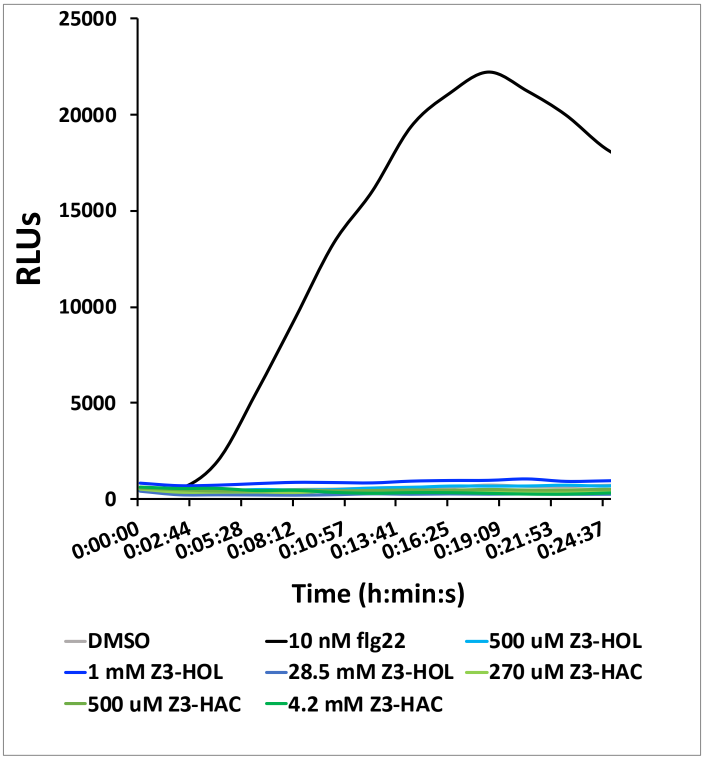


**Supplementary Figure 2. GLVs do not induce an oxidative burst in leaf discs of tomato plants.** Leaf discs were incubated with GLVs and flg22 at the concentrations indicated. ROS were detected with a luminol (L-012)-based bioluminescence assay. Relative light units (RLUs) are shown. No error bars are shown for the GLV experiments (two independent experiments) and DMSO control (four independent experiments) since all responses were close to zero and not statistically different from one another. One representative flg22 experiment is depicted to illustrate classic oxidative burst kinetics.
